# Gas Vesicle-Expressing Human Pluripotent Stem Cells Enable Multimodal Ultrasound and Optical Coherence Tomographic Imaging

**DOI:** 10.1101/2025.04.10.648235

**Authors:** Alessandro R. Howells, John Kim, Sumin Park, Xueding Wang, Chengzhi Shi, Xiaojun Lance Lian

## Abstract

Genetically encoded imaging reporters are critical tools for tracking cell fate and function in regenerative medicine. Gas vesicles (GVs), air-filled protein nanostructures derived from bacteria, offer unique advantages for noninvasive imaging due to their acoustic and optical properties. In this study, we engineered human pluripotent stem cells (hPSCs) to express GVs using a doxycycline-inducible system. Upon doxycycline (Dox) treatment, GVs formed intracellularly and enabled enhanced contrast in both ultrasound and optical coherence tomography (OCT) imaging. Dynamic ultrasound imaging revealed pressure-dependent GV buckling and harmonic signal generation, while OCT imaging confirmed high sensitivity and depth-resolved detection in both *in vitro* and *ex vivo* retinal models. Our work establishes a multimodal GV-based reporter platform compatible with human stem cells and clinically relevant imaging modalities. This approach offers a powerful and versatile tool for noninvasively visualizing and tracking therapeutic cells in real time, advancing the development and monitoring of cell-based therapies.

## INTRODUCTION

Human pluripotent stem cells (hPSCs)^1–3^ hold transformative potential for regenerative medicine due to their capacity for unlimited self-renewal and their ability to differentiate into any somatic cell type^4–10^. These characteristics make hPSCs an ideal source for developing cell-based therapies to repair or replace damaged tissues. However, the clinical translation of hPSC-based therapies is significantly limited by the lack of effective tools for noninvasive real-time monitoring of transplanted cells *in vivo*. Conventional approaches such as immunohistochemistry and histological analysis provide high-resolution insights into cell fate but are inherently endpoint methods that require tissue collection, limiting their utility for long-term studies.

Current *in vivo* imaging strategies often rely on optical techniques such as fluorescence^11^ or bioluminescence imaging. While these methods are useful for visualizing reporter-labeled cells in small animal models, they suffer from limited tissue penetration and resolution at depth. Consequently, their effectiveness diminishes in larger organisms or clinical settings. To overcome these limitations, ultrasound imaging has emerged as an attractive alternative due to its deep tissue penetration, high spatial and temporal resolution, and established clinical accessibility^12^.

Various contrast agents have been explored to enhance cell tracking using different imaging modalities^13–16^. Although exogenous microbubble or nanobubble contrast agents have been employed to track cells using ultrasound, they do require external delivery and are transient in nature, thus limiting their long-term reliability^17,18^. Similarly, nanoparticles have shown promise for cell tracking applications. However, they encounter major challenges in clinical translation due to concerns regarding biocompatibility, clearance pathways, and potential long-term toxicity^19–21^.

A breakthrough in this field was the discovery of genetically encodable ultrasound contrast agents. Unlike exogenous contrast agents that require repeated administration or nanoparticles with uncertain clinical feasibilities, genetically encodable reporters offer several distinct advantages. These reporters enable propagation to daughter cells with each cell division, allowing for long-term monitoring without signal dilution^22^. Furthermore, their genetic nature ensures that only viable cells produce the contrast signal, providing functional information beyond localization^23^.

These genetically encodable contrast agents come in the form of gas vesicles (GVs)^24^. GVs are air-filled protein nanostructures naturally expressed by certain aquatic bacteria and archaea. It’s been shown that GVs enhance ultrasound signals due to their unique acoustic properties^25^. Composed of gas vesicle proteins (Gvps), these nanostructures self-assemble to form intracellular gas-filled compartments that scatter sound waves. Beginning with natural GV-producing organisms, researchers demonstrated that these structures could function as acoustic reporters^25^. Subsequent advances allowed GV expression in *E. coli*^26^ and, more recently, within mammalian cells, giving rise to the concept of mammalian acoustic reporter genes (mARGs)^27^. However, until now, mARG expression has been limited to immortalized or cancer-derived cell lines, and their application to primary or therapeutic stem cells has not been demonstrated.

Here, we present the first successful expression of GVs in hPSCs using second-generation mARGs^28^. The Gvp genes from *Anabaena flos-aquae* can produce 38-fold stronger non-linear acoustic contrast than previously tested clusters when expressed in mammalian cells^28^. We employed a modular gene delivery system combining doxycycline-inducible GvpA expression with constitutive expression of accessory Gvp genes (GvpNtoV), enabling robust and tunable GV production in hPSCs. Microscopy confirmed intracellular GV formation, and multimodal imaging validated their functionality. Specifically, we demonstrated that GV-expressing hPSCs enhance contrast in both static and dynamic ultrasound imaging and are detectable using optical coherence tomography (OCT) *in vitro* and *ex vivo*.

By integrating synthetic biology with noninvasive imaging technologies, our work establishes a new platform for real-time visualization of therapeutic stem cells. This capability has broad implications for regenerative medicine, enabling dynamic tracking of cell fate, biodistribution, and therapeutic efficacy. Our findings position mARGs as a powerful tool for advancing both basic stem cell research and the development of cell-based therapies.

## RESULTS

### Development and microscopy of mARG hPSCs

To engineer hPSCs for GV expression, we utilized the GV gene cluster from *A. flos-aquae*. Specifically, the primary structural protein GvpA was cloned into the doxycycline (Dox)-inducible pXLoneV0 vector (**Fig. 1A**), while the remaining seven secondary genes (GvpN to GvpV) were inserted into a GAPDH donor vector (**Fig. 1A**). This design ensures robust constitutive expression of the secondary genes via knockin downstream of GAPDH, minimizing the likelihood of silencing, and permits precise Dox-dependent induction of GvpA expression through the integrated Tet-On 3G system present in the pXLoneV0 vector. Moreover, the incorporation of distinct drug resistance markers—blasticidin S resistance in the pXLoneV0 construct and hygromycin B resistance in the GAPDH donor vector—facilitates effective selection of the engineered cells.

**Figure 1:**
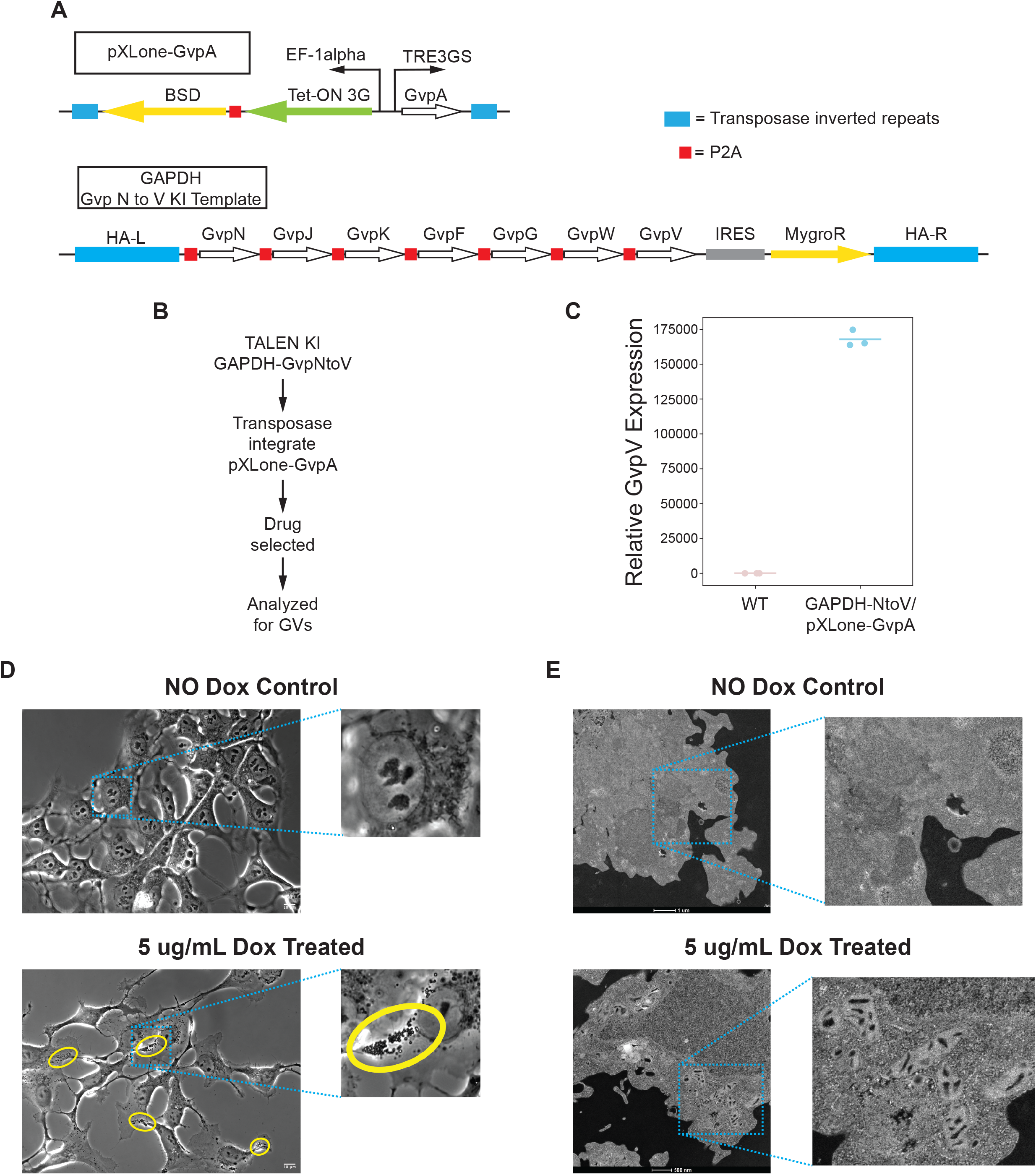
Visualization of GVs within mARG hPSCs. A) Schematic of our two mARG plasmid designs. Top construct is the Dox inducible, transposable pXLoneV0-GvpA cassette, and bottom is the GAPDH-GvpNtoV knock in template. BSD, blasticidin S resistant gene; MygroR, hygromycin B resistant gene; HA-L and HA-R, homology arm left and homology arm right. B) Schematic of workflow to generate our drug selected polyclonal mARG hPSCs. C) qPCR results for relative GvpV expression within wildtype hPSCs and our mARG hPSC line. D) Representative 60x phase contrast images of cells not treated with Dox (top, No Dox Control) and of cells treated with 5 μg/mL Dox (bottom). Yellow ovals indicate where GV aggregates are observed. Scale bars=10 μm. E) Representative electron microscope images of cells not treated with Dox (top, No Dox Control) and of cells treated with 5 μg/mL Dox (bottom). Scale bars=1 μm and 500 nm.

Following GAPDH knock-in and transposase-mediated integration of pXLoneV0-GvpA into H1 hPSCs, we selected a polyclonal population using the corresponding antibiotics (**Fig. 1B**). Validation by quantitative PCR (qPCR) revealed that relative GvpV RNA levels in the engineered cells were approximately 170,000-fold higher than those in wildtype H1 cells, confirming successful integration and selection (**Fig. 1C**).

We further corroborated GV formation using microscopy. Phase contrast imaging at 60x magnification of cells treated with 5□μg/mL Dox for 72 hours revealed cytoplasmic structures indicative of GV aggregates, in contrast to the clear cytoplasms observed in untreated control cells (**Fig. 1D**). Additionally, electron microscopy demonstrated that while control cells maintained normal cytoplasmic appearance, Dox-treated cells exhibited distinct, gas-filled vesicular structures, likely representing cross sections of the GVs (**Fig. 1E**).

### Gas vesicles as a potential contrast agent for static ultrasound imaging

To demonstrate the feasibility of using GVs as contrast agents in ultrasound imaging, we used our established mARG H1 hPSCs prepared with or without Dox induction, alongside wild-type H1 hPSCs to serve as a control. We first performed simplified static imaging, without high intensity ultrasound, to induce GV buckling as previously described^27,29^ (**Fig. 2A**). We hypothesized that a concentration of GV-containing cells would scatter more ultrasound signal as a cluster due to the acoustic impedance mismatch between the gas content in GVs and the fluids in the cytoplasms, even when the vesicles are sub-wavelength in size.

**Figure 2:**
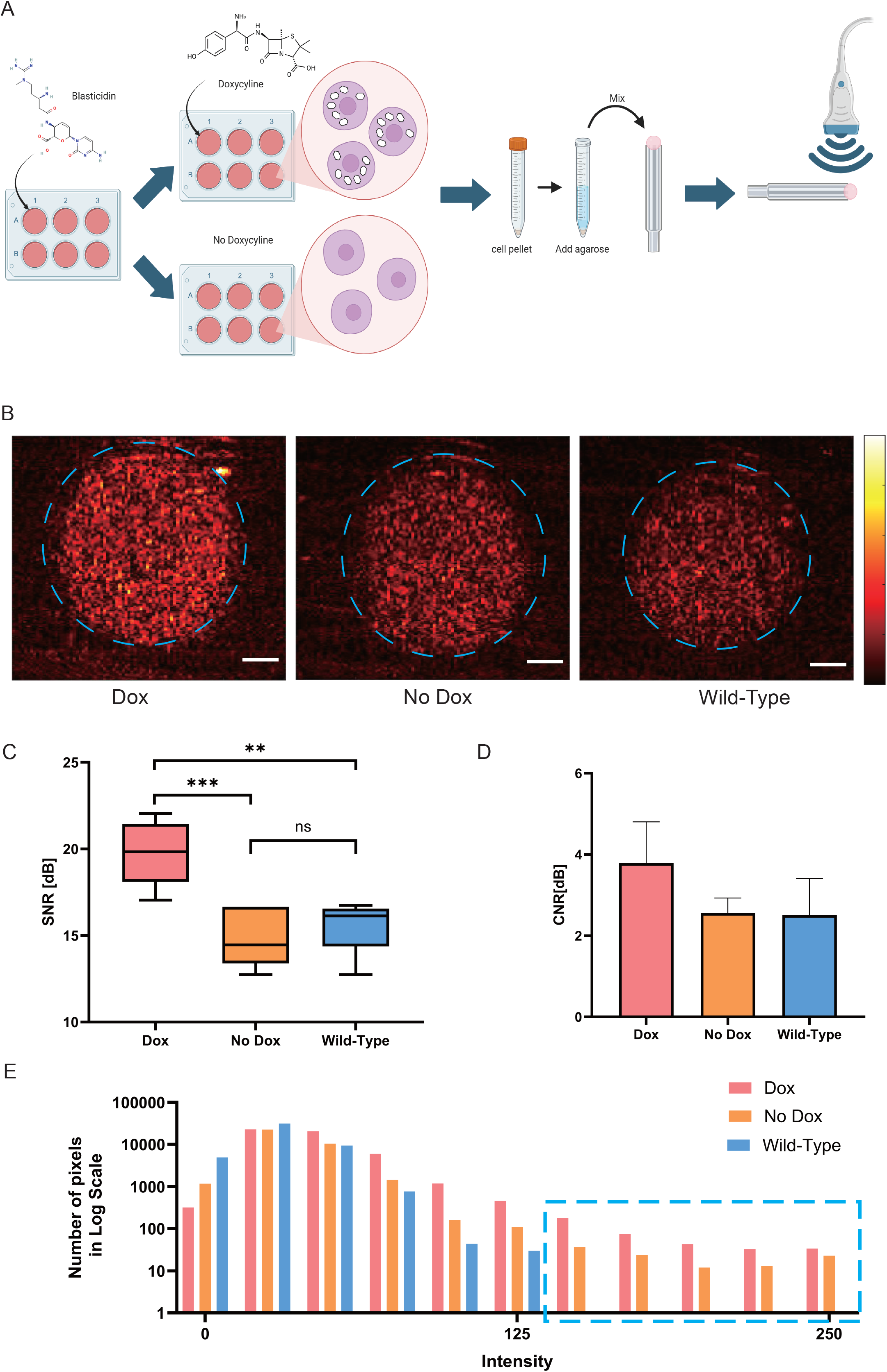
Static ultrasound images of GV-expressing hPSCs. A) schematic drawing of static imaging set up that shows from drug selectable GV expressions on cells to imaging. B) Ultrasound cross section images of cell samples mixed with 1% agarose gel. The hot-scale B-mode images present the Dox treated mARG hPSCs (left), the non-doxycycline treated mARG hPSCs (middle), and wild-type stem cells (right). Color scale: 1e3–1e5. C) SNR values for different conditions in decibels (dB). The Dox-treated samples exhibit significantly higher SNR compared to No Dox and Wild-type groups (**p□< □0.01. ***p□< □0.001, ns: not significant). D) CNR values in dB for each condition. The Dox-treated samples demonstrate the highest CNR, indicating enhanced contrast between the gas vesicle-expressing cells and the background. E) Histogram (log scale) represents the distribution of pixel intensities. Dox-treated samples exhibit a higher number of high-intensity pixels compared to No Dox and Wild-type samples, as highlighted in the dashed blue box. Scale bars=1 mm.

Ultrasound imaging of cell clusters with Dox-treatment revealed pronounced contrast in B-mode imaging, observable even prior to background subtraction (**Fig. 2B**). Quantitative analysis of the signal-to-noise ratio (SNR) indicated a marked difference between wild-type and Dox-treated cells. The SNR, quantified for each condition, demonstrated that Dox-treated mARG hPSCs exhibited a significantly higher SNR (19.7 ± 1.9 dB) compared to the untreated (14.9 ± 1.7 dB) and wild-type (15.6 ± 1.6 dB) samples, thereby confirming that GV expression contributes to enhanced ultrasound signal (**Fig. 2C**). The minimal difference between untreated and wild-type cells further substantiates that in the absence of Dox, GV expression does not occur.

Additionally, contrast-to-noise ratio (CNR) values were calculated to assess the delineation of the GV–expressing region from the surrounding background. As expected, the Dox-treated mARG hPSCs exhibited the highest CNR (3.8 ± 1.0 dB). However, the differences among the groups (Dox, No Dox, Wildtype) were less pronounced compared to those observed in the SNR analysis, suggesting that while GV expression improves ultrasound contrast, the absolute signal strength (as evaluated by SNR) serves as a more robust indicator of GV presence (**Fig. 2D**). A histogram on a logarithmic scale of pixel intensity distribution across the different treatment conditions further highlighted the presence of high-intensity pixels in the Dox-treated samples (**Fig. 2E**). Specifically, the highlighted blue box in the histogram indicates a greater number of high-intensity pixels in the Dox-treated group, consistent with enhanced signal output from GVs, while untreated and wild-type samples demonstrated a lower and more uniform intensity distribution (**Fig. 2E**).

### Dynamic imaging of GVs in mARG hPSCs

We previously demonstrated that simple drug selection can effectively select a mixed population of GV expressing HEK293 cells^29^. We showed this by using focused ultrasound (FUS) at a mechanical index (MI, a measure of the potential of acoustic cavitation and the corresponding damage to tissues) lower than the FDA regulation of 1.9^30^. At this lower level, we were therefore able to induce GV buckling without causing any acoustic damage to the cells. However, with the combination of synchronized FUS induced GV buckling and a B-mode imaging system, we can capture dynamic ultrasound contrast.

Building upon the previous static ultrasound imaging setup, FUS was applied in a vertical orientation to facilitate simultaneous GV buckling and real-time imaging, which was optimized for detection, sensitivity, and spatial resolution (**Fig. 3A**). Unlike the static imaging results, post-processing was applied to the images to delineate the outcomes following FUS exposure (**Fig. S1A**). Post-FUS frames were subtracted from pre-FUS frames to analyze the GV dynamic contrast induced by the FUS. The input peak negative pressures (PNP) ranged from 0.5 to 3 MPa and all corresponded to a mechanical index of less than 1.2 (**Fig. S1B**).

**Figure 3:**
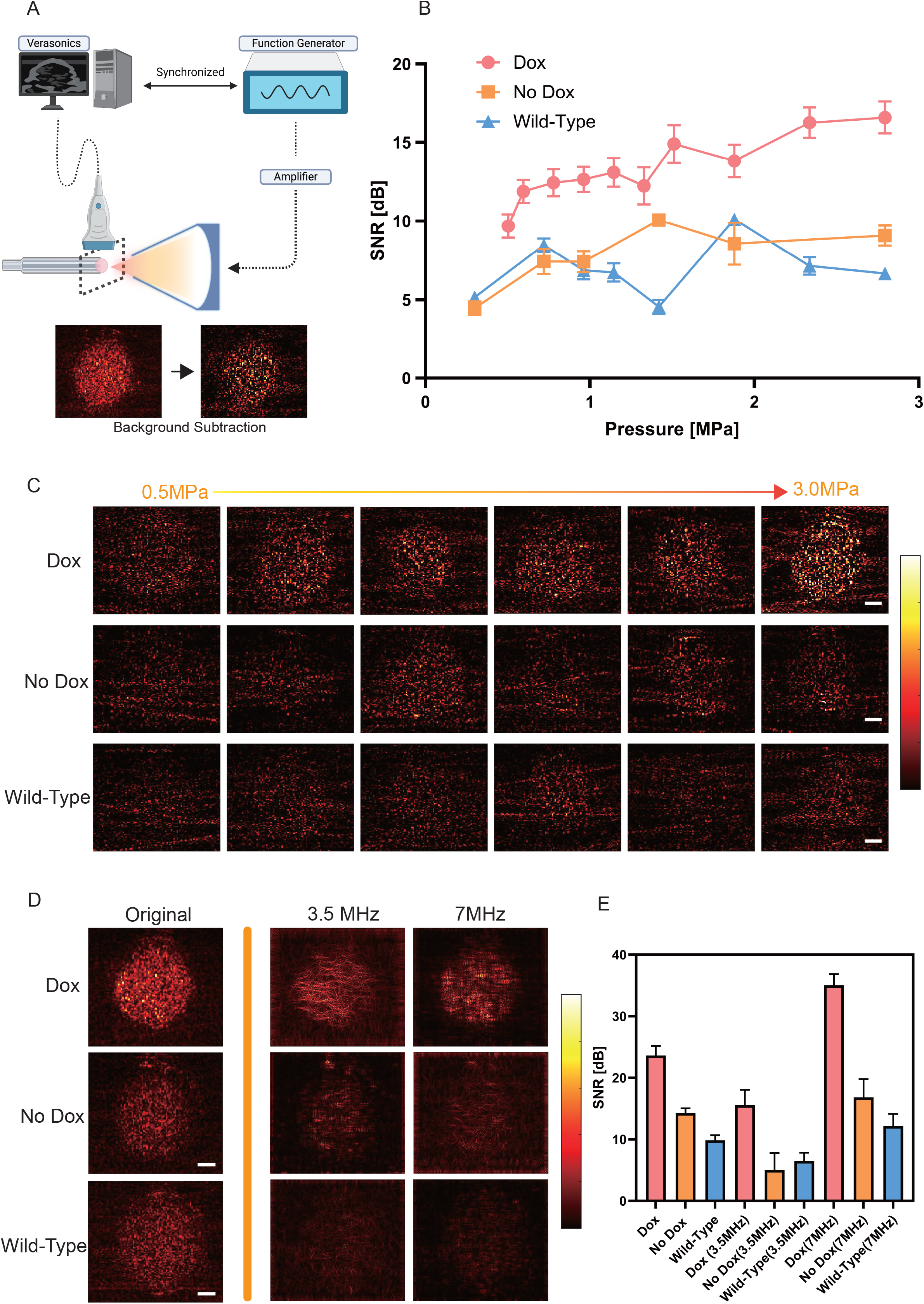
Dynamic ultrasound images of GV-expressing hPSCs. A) The schematic illustrates the experimental setup for FUS stimulation and subsequent imaging that are synchronized. Ultrasound response was analyzed after background subtraction to enhance signal detection. B) The SNR values of the region of interest (ROI) are plotted against increasing ultrasound pressure (0–3 MPa) for three sample conditions (Dox, No Dox, Wild-type). Dox-treated samples exhibit a continuous increase in SNR with increasing pressure, whereas No Dox and Wild-type samples show significantly lower and more variable SNR responses. C) A series of ultrasound images demonstrates the effect of increasing FUS pressure (0.5 MPa to 3.0 MPa) on different sample conditions. Dox-treated samples show a progressive enhancement in brightness and signal intensity, particularly at higher pressures, suggesting GV buckling. Color scale: 1e3–1e5. D) Ultrasound images were processed with frequency-based filtering (3.5 MHz and 7 MHz) to enhance harmonic signal detection. Dox-treated samples show distinct harmonic components at 3.5 MHz and 7 MHz, while No Dox and Wild-type samples display weak or no discernible harmonic signals. Color scale: (Original)1e3–1e5/ (3.5 and 7MHz): 1e3–3e4. E) The bar graph quantifies SNR values for the original and frequency-filtered images. Dox-treated samples exhibit significantly higher SNR across all conditions, particularly in the 3.5 MHz and 7 MHz filtered images, reinforcing the role of gas vesicles in enhancing ultrasound contrast. No Dox and Wild-type samples maintain consistently lower SNR values, indicating the absence of detectable gas vesicle activity. Scale bars=1 mm.

The SNR of the Dox-treated mARG hPSCs steadily increased as the pressure increased, eventually plateauing at higher levels (**Fig. 3B**). This behavior indicates that GV buckling can enhance the acoustic response, but is limited by the concentration of GVs. In contrast, both the No Dox and Wild-type cells maintained significantly lower SNR values throughout all pressure ranges tested. Notably, the divergence in SNR between the Dox-treated and control groups becomes pronounced at pressures above 1.5 MPa, suggesting that GV-containing cells are more susceptible to experience acoustic cavitation, potentially accompanied with cellular damage^31,32^.

These results confirm that GV expression markedly enhances ultrasound contrast in a pressure-dependent manner, while non-expressing cells remain relatively unaffected (**Fig. 3B,3C**). Ultrasound images were processed using frequency-based filtering at 3.5 MHz and 7 MHz to boost harmonic signal detection. Dox-treated mARG hPSCs exhibit distinct harmonic components at both frequencies, whereas the No Dox and wild-type samples show weak or undetectable harmonic signals, confirming that the enhanced nonlinear acoustic signal arises from GV activity (**Fig. 3D,3E**). Additionally, the bar graph quantifies SNR values for both the original and filtered images. Dox-treated samples consistently displayed significantly higher SNRs (23.6 ± 1.5 dB), especially in the frequency-filtered images (3.5MHz: 15.6 ± 2.4 dB, 7MHz: 35.1 ± 1.8 dB), while No Dox and Wild-type samples maintained lower SNR values, underscoring the absence of detectable GV activity.

### *In vitro* and *ex vivo* OCT imaging of GVs in mARG hPSCs

GVs can be used as OCT contrast agents. To evaluate whether GVs can be detected via OCT when they were expressed in hPSCs, a custom-modified SD-OCT system was developed for phantom and ocular imaging, as illustrated in **Fig. 4A**. This system demonstrated high sensitivity and depth-resolved imaging capabilities. Four agarose phantom groups embedded with Dox-treated mARG hPSCs were fabricated at concentrations of (i) 5 ×□10^6^ cells/mL, (ii) 1 ×□10^6^ cells/mL, (iii) 5 ×□10^5^ cells/mL, and (iv) 1 ×□10^5^ cells/mL.

**Figure 4.**
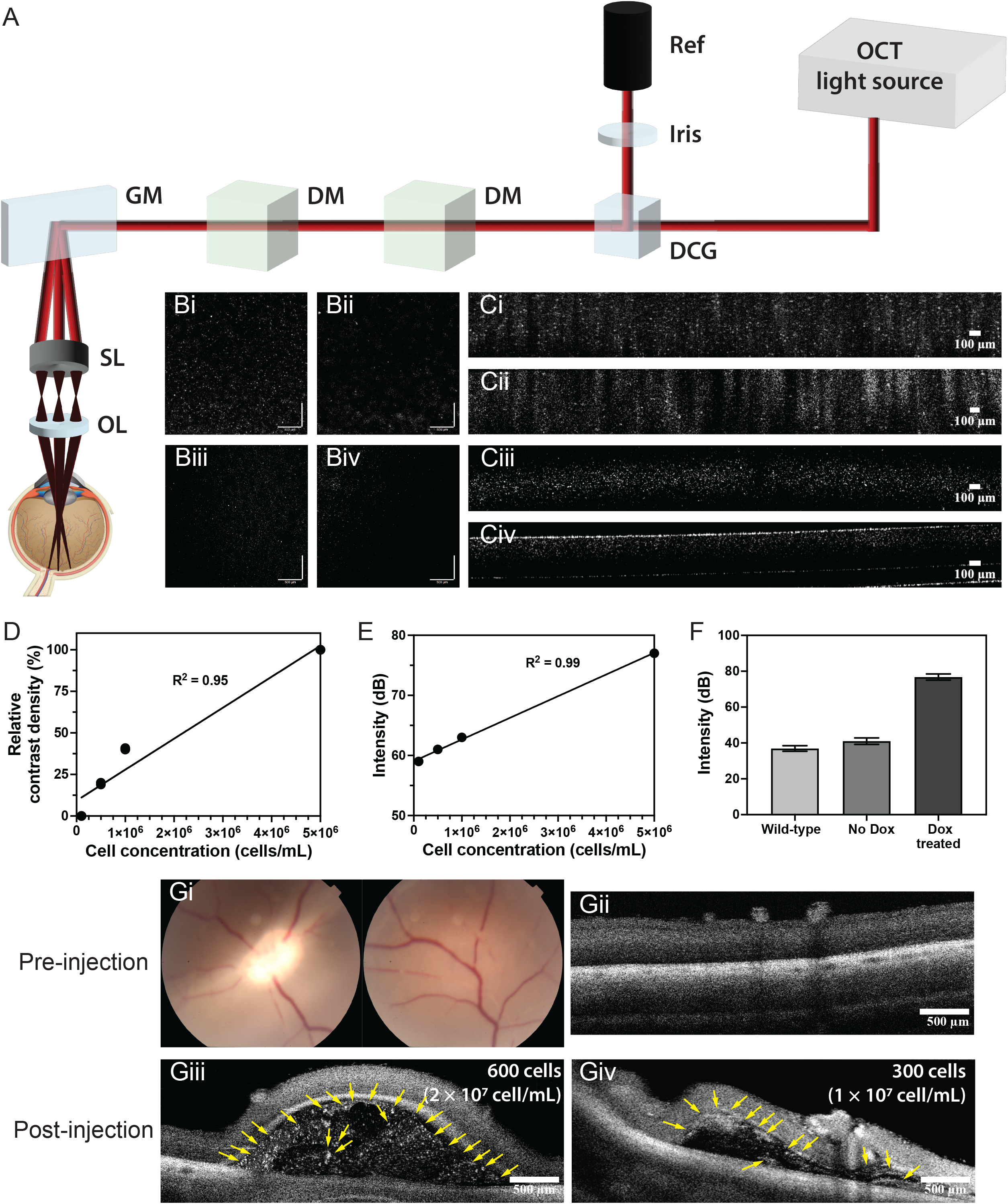
OCT of GV-expressing stem cells. A) Schematic diagram of the OCT system used for imaging GV-expressing hPSCs. (Ref reference arm; DCG dispersion compensation glass; DM dichroic mirror; GM galvanometer; SL scan lens; OL ophthalmic lens). Representative OCT images of agarose phantoms with GV-expressing hPSCs: Bi – Biv) en-face images and Ci – Civ) B-scan images at concentrations of (i) 5 ×□10^6^ cells/mL, (ii) 1 ×□10^6^ cells/mL, (iii) 5 ×□10^5^ cells/mL, and (iv) 1 ×□10^5^ cells/mL, respectively. Quantitative analysis: D) relative signal density versus cell concentration (R^2^ = 0.95), and E) signal intensity versus cell concentration (R^2^ = 0.99). F) Intensity comparison among wild-type, non-DOX-treated, and DOX-treated in vitro samples. G) Ex vivo imaging of porcine eye: Gi) Fundus photograph showing a healthy optic disc and retinal vasculature before injection. Gii) Cross-sectional OCT image of the porcine retina before injection. OCT image post-injection of Giii) 600 cells at a concentration of 2 × 10^7^ cells/mL, and Giv) 300 cells at a concentration of 1 × 10^7^ cells/mL. The yellow arrows indicate gas vesicle-expressed individual stem cells.

OCT images of the phantoms revealed uniform cell distribution within the agarose matrix. En-face images (**Fig. 4Bi – Biv**) showed higher signal densities with increasing cellular concentrations, while B-scan images (**Fig. 4Ci – Civ**) provided depth-resolved information. These results confirmed the sensitivity of the system, revealing enhanced contrast by the GV expressing mARG hPSCs. These findings are significant for establishing the feasibility of this imaging approach in preclinical studies requiring cellular visualization and localization. The statistical analysis shown in **Fig. 4D-E** confirmed a linear correlation between cell concentration and both signal density (R^2^ = 0.95) and peak intensity (R^2^ = 0.99). This is evidenced by the statistical analysis of contrast density and signal amplitude, which shows a significant signal density and intensity increase with higher cell concentrations. In addition, to confirm the observed signals were generated from the GVs, agarose phantoms were embedded with wild-type hPSCs and non-DOX-treated mARG hPSCs. The statistical analysis is shown in **Fig. 4F**. WT (36.9 ± 1.5 dB) and non-DOX-treated (41.0 ± 1.8 dB) samples showed significantly lower signal intensities compared to the DOX-treated sample (76.8 ± 1.7 dB). The DOX-treated sample indicated around 1.8 – 2 times higher signal intensities than other samples.

By confirming the reliable and high-resolution imaging system for GV-expressing stem cells from the above *in vitro* study, we set up an *ex vivo* experiment to test the feasibility of monitoring mARG hPSCs within ophthalmic tissue. **Fig. 4G** presents fundus photographs and OCT images of a porcine eyeball acquired *ex vivo*. The fundus photograph (**Fig. 4Gi**) displays a clear view of the optic disc and retinal vasculature prior to injection, indicating a healthy baseline condition. The absence of morphological abnormalities confirms the structural integrity of the sample before the injection. A baseline cross-sectional OCT image of the retina (**Fig. 4Gii**) further establishes a reference for subsequent imaging comparisons. Following the subretinal injection of GV-expressing stem cells at varying concentrations, OCT images revealed localized contrast signals within the subretinal space. At a concentration of 2 × 10^7^ cells/mL (**Fig. 4Giii**) and 1 × 10^7^ cells/mL (**Fig. 4Giv**), the injected cells produced distinct hyper-reflective signals. Quantitative analysis of OCT signal intensity demonstrated that the retinal layer exhibited a mean intensity of 46.6 ± 2.2 dB, while the GV-expressing stem cells generated a significantly higher intensity of 59.8 ± 2.3 dB. This corresponds to a 128% enhancement in signal contrast compared to the surrounding ophthalmic tissue, confirming the detectability of mARG hPSCs within the retinal microenvironment.

These results highlight the feasibility of imaging and tracking GV-expressing stem cells by OCT, underscoring their potential applications in tissue engineering and regenerative medicine including those in ophthalmology clinics.

## Discussion

In this study, we demonstrate the successful engineering of hPSCs to express GVs for multimodal imaging applications. By leveraging a doxycycline-inducible system for GvpA expression and constitutive expression of the remaining GV structural genes, we achieved robust and controllable GV production in hPSCs. Microscopy data confirmed the presence of intracellular gas-filled vesicles upon induction, validating our genetic design. Importantly, the use of dual antibiotic selection helped ensure integration fidelity and enabled efficient enrichment of GV-positive cells, offering a streamlined workflow for producing reporter stem cell lines.

The ability of GV-expressing hPSCs to enhance contrast in static and dynamic ultrasound imaging was clearly demonstrated. Cells with Dox treatment produced significantly higher SNR and CNR ratios compared to controls, indicating successful acoustic modulation by intracellular GVs. Under FUS stimulation, we observed pressure-dependent buckling behavior, a hallmark of functional GV activity, as well as harmonic signal generation. Importantly, the observed pressure-dependent phase transitions in GVs enabled both enhanced imaging contrast and the potential for cavitation-mediated modulation, expanding their functional utility beyond passive contrast enhancement. Notably, these dynamic responses were absent in wild-type and cells without Dox treatment, reinforcing the specificity and reliability of GV expression as an imaging marker.

We further extended our findings by applying OCT, where GV-expressing stem cells showed strong signal intensities both *in vitro* and in *ex vivo* porcine retina models. Quantitative analyses demonstrated a clear correlation between cell concentration and OCT signal strength, establishing the sensitivity of this imaging platform. In the *ex vivo* retinal setting, injected cells remained detectable with enhanced contrast relative to the surrounding tissue, albeit at slightly reduced levels compared to *in vitro* results. These data suggest that while GVs are effective optical scatterers, physiological factors in tissue may attenuate signal propagation and highlight the need for further optimization under *in vivo* conditions. To further enhance the sensitivity of OCT in imaging the GV-expressing stem cells in ophthalmic tissues, increase the optical scattering contrast generated by the GVs in these cells will be a key focus of our future research. Such advancements would establish GV-expressing stem cells as a platform tool for understanding cell-based therapies in preclinical and clinical settings in ophthalmology.

Collectively, our work establishes GVs as genetically encoded, multimodal contrast agents compatible with ultrasound and OCT imaging in stem cells. This platform holds significant promise for noninvasive tracking of therapeutic cells in regenerative medicine and real-time monitoring of cell-based therapies. Future work will focus on optimizing GV expression to enhance signal robustness *in vivo*, evaluating long-term stability of the reporter system, and extending this strategy to additional cell types and disease models. Integration of GV-based imaging into clinical workflows could transform our ability to monitor cellular dynamics with high spatial and temporal resolution.

## Material and Method

### Plasmid cloning

All plasmids were cloned using the Takara In-Fusion cloning kit. Primers were designed by the Takara In-Fusion online portal. For pXLoneV0-GvpA, XLone-GFP (Addgene # 96930) was restriction enzyme digested with SpeI and KpnI. The GvpA insert was PCR amplified from a plasmid gifted to us from Dr. Mikhail Shapiro. For the GAPDH-GvpNtoV plasmid, pUC-GW-Amp backbone was digested with MluI and AscI. GAPDH homology arms and IRES-MygroR were PCR amplified from pGAPTrap-mCherry-IRESMygro (Addgene # 82505) and GvpNtoV were PCR amplified from a plasmid gifted to us from Dr. Mikhail Shapiro. After 10 μL In-Fusion reaction, 5 μL were transformed into Stbl3 Chemically Competent E. coli following manufacturing protocol. E.coli were plated onto Ampicillin selection plates and cultured overnight. Clones were then picked and cultured for 18 hours in LB broth supplemented with Ampicillin. Finally, plasmids were harvested via Zymopure plasmid miniprep kit and sequenced.

### hPSC maintenance and genetic engineering

hPSCs were cultured on iMatrix-511 coated Falcon well plates, in either essential 8 or mTeSR1 media. Routine passages were performed once confluency reached about 70% by dissociating with accutase and centrifuging for 4 minutes at 200 RCF. Cell pellets were then resuspended in media containing ROCK inhibitor and plated onto freshly coated wells.

For TALEN mediated knock in of our GAPDH-GvpNtoV construct, wildtype H1 hPSCs were pelleted and resuspended in 100 μL PBS buffer containing 10 μg of our donor vector, 5 μg of pGAPTrap-TALEN 1 (Addgene #83368), and 5 μg of pGAPTrap-TALEN 2 (Addgene #83369). Cells were then transferred to a cuvette and nucleofected on program CB150 on a Lonza 4D Nucleofector. Cells were then plated at high density and after expansion, were drug selected to a final concentration of 200 μg/mL hygromycin B. For pXLoneV0-GvpA integration, the same nucleofection protocol was followed, however they were nucleofected with 5 μg pXLoneV0-GvpA and 5 μg EF1a-hyPBase transposase plasmids, and were drug selected to a final concentration of 20 μg/mL BSD.

### Quantitative Polymerase Chain Reaction (qPCR)

RNA was collected from hPSC samples using the Direct-zol RNA MicroPrep kit (Zymo Research) according to the manufacture’s instructions. cDNA synthesis was then carried out on 500 ng RNA using the ZymoScript RT Premix (Zymo Research). cDNA was then diluted to 2 ng/μL. qPCR reactions were then set up in triplicate with 10 μL Power SYBR Green Master Mix (ThermoFisher), 1 μL cDNA (2ng/μL stock), 8 μL molecular biology grade water, and 1 μL primer mix (5 μM stock). Forward GvpV primer used was TGTACAAGATGGTCACCGAGAA and reverse GvpV primer was TGTTGCTTTCCACGAAGATGTT. Housekeeping gene used for normalization was GAPDH, (forward: GTGGACCTGACCTGCCGTCT and reverse: GGAGGAGTGGGTGTCGCTGT). qPCR reactions were then run on the CFX Connect Real-Time System (Bio Rad) with an initial 10-minute incubation at 95°C and 40 cycles of 15 seconds at 95°C followed by 75 seconds at 60°C. Relative GvpV expression was then calculated based on Cq values and normalized to GAPDH.

### Phase contrast imaging and electron microscopy

Phase contrast was performed by plating cells sparsely onto glass bottom wells. Cells were then cultured for a total of 72 hours, either with or without 5 μg/mL Dox, and the day prior to imaging, media was changed containing ROCK inhibitor. Phase contrast images were acquired using a custom-built microscope based on an Olympus IX-73 body, equipped with Olympus phase contrast objectives, a Kiralux 12.3 MP monochrome CMOS camera (CS126MU, Thorlabs), a 100W halogen lamp (Olympus), and a SOLA U-nIR light engine (Lumencor).

For electron microscopy, about 1 million cells were fixed with 2.5% glutaraldehyde/4% Paraformaldehyde and post fixed with 1% Osmium Tetroxide in 0.1M Sodium Cacodylate Buffer. Sample were then stained with uranyl acetate, dehydrated with increasing concentrations of ethanol, and embedded in Epon Epoxy. Grids were then imaged using scanning electron microscopy and transmission electron microscopy (S/TEM Talos F200×2 from Thermo Scientific).

### Sample preparation for ultrasound imaging

Two 6-well plates were prepared for each experiment to acquire samples for ultrasound imaging. Before ultrasound experiment, stem cells were treated with doxycycline for 72 hours to assure abundant time for GVs to fully grow. The cells were dissociated with Accutase, and counted with Cell counting device (Cell Counter 3 FL, ThermoFisher) at a concentration of 20 million cells/ml. The harvested cells were centrifuged and resuspended in 1% agarose gel (ThermoFisher) to create a solidified sample within a plastic holder, ensuring a stationary position in a water tank filled with deionized, degassed water at 37°C. A volume of 200 μl of the cell-gel mixture was layered on top of the holder, allowing the cells to not only fill the holder but also form an additional layer above it. Such method allowed optimal transducer alignment, minimizing imaging of the outer perimeter of the plastic sample that could otherwise introduce sound artifacts.

#### Imaging transducer

For ultrasound imaging, vantage NXT was used to collect data with transducer L22-8v. The position of the transducer was fitted vertically upward from the sample that it can cover cross section area of the mixed cell sample. The center frequency was set to 15.6MHz that is covered with 200% Nyquist frequency. For ultrasound imaging acquisition, three angles with three apertures were utilized to generate each frame, resulting in a total of nine transmit-receive pairs per image. The time between each event were 200us which operates approximately at 800 frames per second.

#### Focused ultrasound

A single-element focused ultrasound transducer operating at 3.5 MHz (H-101, SonicConcepts) was fitted with a coupling cone backfilled with degassed, deionized water such that the focus point is at the tip of the coupling cone. The structural input of focused ultrasound where it is synchronized to Verasonics, sending out signals and amplified at 3.5 MHz. Focused ultrasound transducer was connected to both function generator and amplitude so that it sends out 10 cycles of desired input voltage only when triggered by imaging system. The synchronized set up assured the repeatability and consistence of experimental outcome

#### Post Processing

Post-processing was performed to enhance signal detection, quantify ultrasound contrast, and evaluate gas vesicle (GV) activity under focused ultrasound (FUS) stimulation. Ten pre-collapse frames and forty post-collapse frames were captured per 10-cycle burst sine wave. Ten pre-frames were averaged to be subtracted from the post frames images. Among the subtracted images, SNR and CNR was collected from ROI. ROI was defined as same as static image.

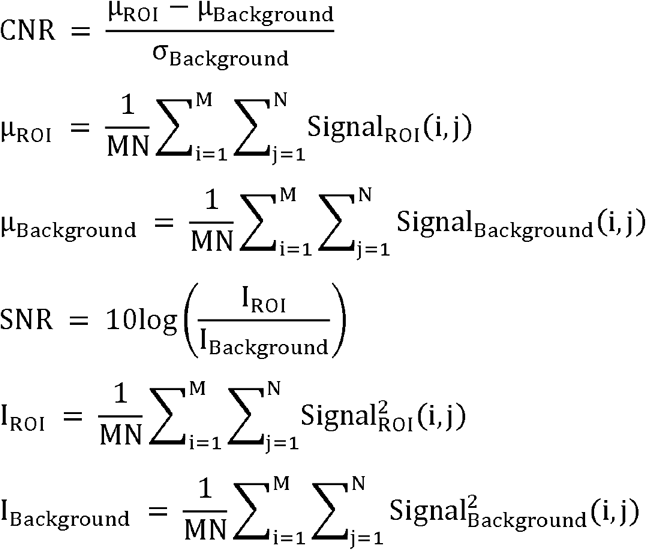

In these equations, M and N are the dimensions of the ROI and background in pixels, *Signal*_*ROI*_ (*i, j*) and *Signal*_*Background*_ (*i, j*) are the signal intensity at pixel (*i, j*) within the ROI and background area, respectively. *I*_*ROI*_ is the average signal intensity of the ROI, *I*_*Background*_ is the average background signal intensity.

### Sample Preparation for OCT imaging

#### Agarose phantom preparation

To mimic the optical and mechanical properties of biological tissues and ensure reproducible and controlled experimental conditions, we used a 1% w/v agarose gel (Sigma Aldrich, St. Louis, MO, USA) in this study^33,34^. Agarose powder was dissolved in deionized water by heating to 90 °C under constant stirring until fully homogenized. The solution was degassed and cooled to 37 °C to ensure uniformity and minimize bubble formation. GV-expressing cells were centrifuged, resuspended in the agarose solution at a predefined concentration, and cast into a customized mold. The gel solidified within minutes. The optical transparency and acoustic impedance of the agarose phantom closely resembled those of biological soft tissues, making it suitable for calibration and validation of imaging systems.

#### Porcine eyeball preparation

The use of porcine eyes as surrogates for human tissues is well-established due to their anatomical and physiological similarities^35^. Fresh porcine eyeballs were obtained from a local slaughterhouse, where all animals were euthanized on the same day as the study to preserve tissue hydration and structural integrity. The eyeballs were transported at 4 °C in Dulbecco’s Modified Eagle’s Medium (DMEM; Gibco, Waltham, MA, USA) supplemented with 1% penicillin and streptomycin (ThermoFisher), 1% L-glutamine (Gibco), 10% fetal bovine serum (FBS, Corning, NY, USA) to maintain tissue viability and prevent contamination^36^. Excess connective tissues were removed, and the eyeballs were disinfected using a 1:2 (v/v) solution of povidone-iodine and phosphate buffered saline (PBS, Gibco, Waltham, MA, USA) for 5 minutes. Following disinfection, the eyes were rinsed with sterile PBS.

A 30 μL of GV-expressing cell mixture at different concentrations (2 × 10^7^ and 1 × 10^7^ cells/mL) was injected into the subretinal space. The subretinal injection was performed following a published protocol^37^. Briefly, the eyeball was secured in a customized holder under a dissecting microscope. The superior rectus muscle was excised using sterile scissors, and a scleral tunnel was created 3.5 mm posterior to the limbus using a 26-gage sterile disposable needle. A contact lens coated with Gonak gel (Akorn Inc., Lake Forest, IL, USA) was placed on the cornea to stabilize the ocular surface and visualize the injection site. Using a 30-gauge Hamilton syringe loaded with the cell suspension, the needle was carefully introduced through the scleral tunnel and advanced under microscopic guidance until it approached the retinal space. The cell suspension was slowly injected with care to avoid disruption to the retinal tissue. After the injection, the syringe was gradually retracted to minimize the risk of tissue damage.

#### Imaging Systems

For *in vitro* and *ex vivo* imaging, a spectral domain optical coherence tomography (SD-OCT, TEL321, Thorlabs, Newton, NJ, USA) system modified with an ophthalmic lens (AC080-010-A) was used^38,39^. SD-OCT is a reliable and non-invasive imaging technique that employs infrared light to detect backscattered photons by low-coherence interferometry, allowing high-resolution analysis of the anatomical structures of the eye.

The OCT system employed an illumination light source with a center wavelength of 1300 nm. The light beam was split into reference and sample arms. In the reference arm, reflected light was directed back into the light source fiber, with the intensity of the reference light adjusted using an iris. In the sample arm, the beam passed through a 2D galvanometer and was focused onto the sample using a scan lens (LSM03, Thorlabs) and ophthalmic lens. The light reflected from the sample was combined with the reference light in the interferometer to produce interference patterns, which were detected by a line scan camera operating at up to 146 kHz repetition rates. The system achieved lateral and axial resolution of 13.0 µm and 5.5 µm in air, respectively. For all OCT images, A-lines acquisition was conducted at a rate of 10 kHz with a scanning area of 1024 × 400 points, providing an axial field of view of 1 mm.

#### Data analysis

The OCT data were acquired through ThorImageOCT software (Thorlabs), and images were processed with MATLAB (Mathworks Inc., Natick, MA, USA) and ImageJ (National Institutes of Health, Bethesda, Maryland, USA). The statistical analysis was completed with GraphPad Prism 10 (GraphPad Software Inc., San Diego, CA, USA).

## Acknowledgments

This work was supported by NSF CBET-1943696 (to X.L.L.), CBET-2319913 (to X.L.L.), NIH R56DK133147 (to X.L.L.), R01HL175050 (to X.L.L.), R01EY034325 (to X.W.), Huck Innovative and Transformational Seed (HITS) Fund (to X.L.L.). This work was also supported by NSF CMMI-2142555 (to C. S.). We thank Dr. Pentao Liu’s lab for kindly sharing their EF1a-hyPBase plasmid. The data sets obtained and used in this study are available upon request submitted to the corresponding author.

## Author Contributions

A.R.H, J.K., and S.P. performed experiments, analyzed data, and wrote the manuscript. C.S. and X.W. analyzed data, wrote the manuscript, and supervised the project. X.L.L. performed experiments, analyzed data, wrote the manuscript, and supervised the project.

## Figure Legends

**Figure S1.** A) Schematic representation of the experimental protocol for frame acquisition before and after focused ultrasound (FUS) exposure. Agarose mixed cells were imaged using high-frame-rate ultrasound imaging before and after the application of a single FUS pulse. The FUS was delivered at the center of the imaging region. Pre-FUS frames were acquired immediately prior to FUS exposure, while post-FUS frames were acquired following the pulse. The highlighted regions represent frames used for pre-FUS and post-FUS analysis. B) Mechanical index (MI) as a function of peak negative pressure used during FUS stimulation. This linear relationship illustrates the range of mechanical indices corresponding to the pressures applied during the experiment (0–5 MPa. The MI was calculated using the standard formula where peak negative pressure in MPa, and f is the transmit center frequency in MHz (assumed to be 3.05 MHz in this study), MI=P_neg_/sqrt(f).

