## Supplementary figures and images for "Gas Vesicle-Expressing Human Pluripotent Stem Cells Enable Multimodal Ultrasound and Optical Coherence Tomographic Imaging"

### Supp Fig 1

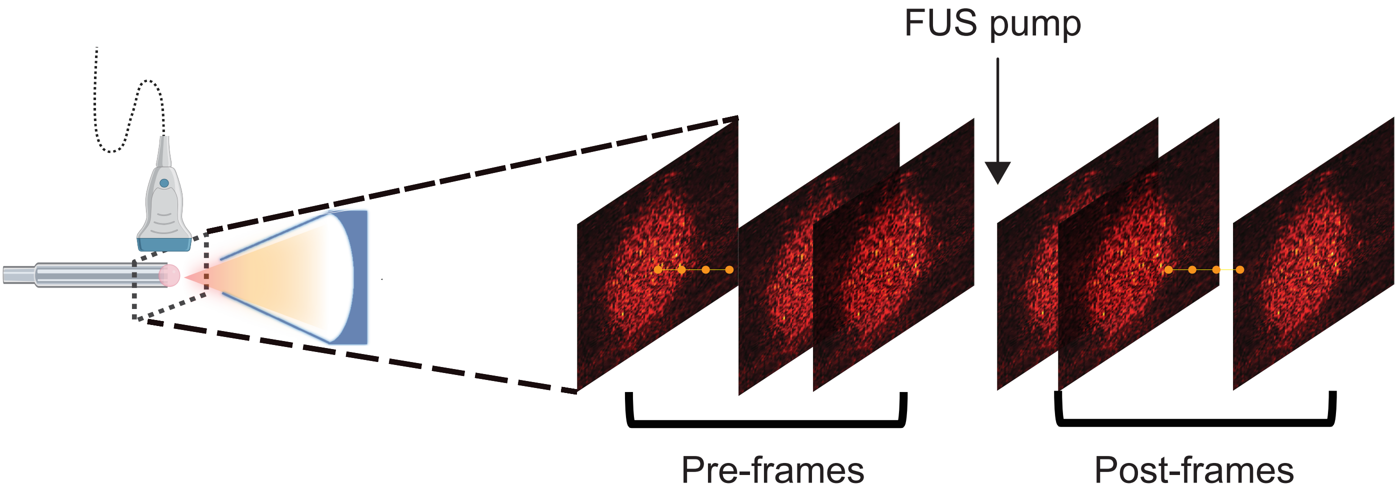

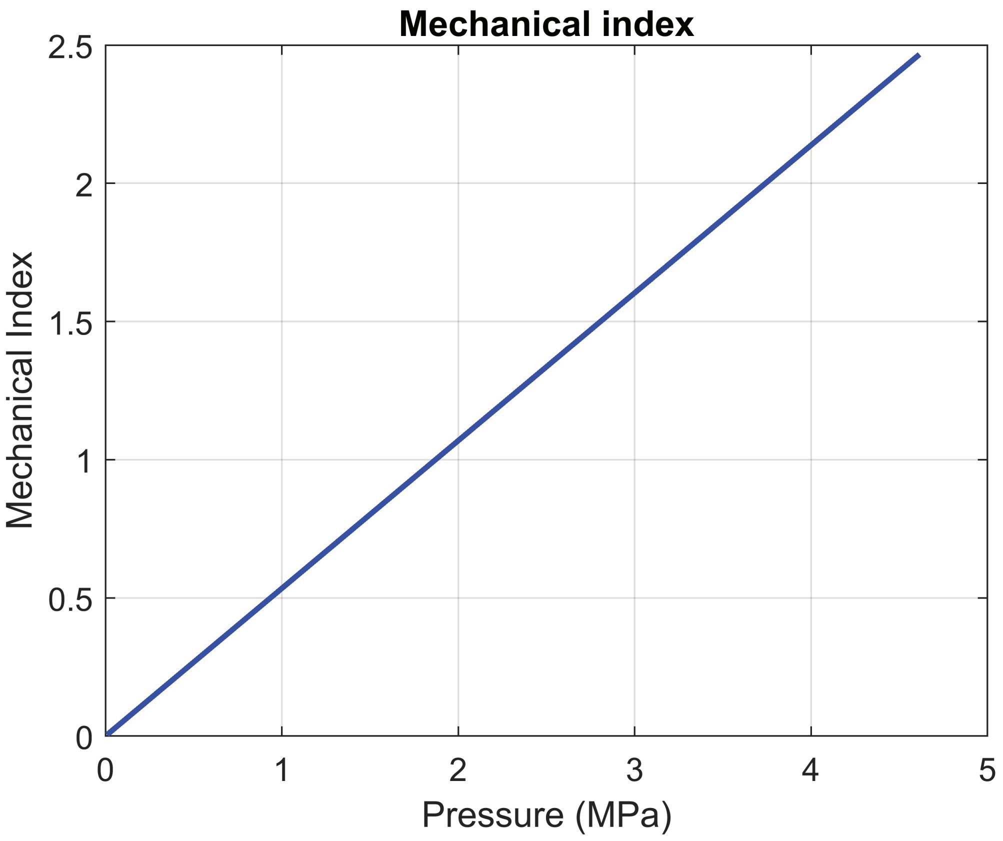
